# Ontogenesis of the tear drainage system requires Prickle 1-controlled polarized basement membrane (BM) deposition

**DOI:** 10.1101/2020.04.07.029793

**Authors:** Dianlei Guo, Jiali Ru, Fuxiang Mao, Kaili Wu, Hong Ouyang, Yizhi Liu, Chunqiao Liu

## Abstract

In terrestrial animals, lacrimal drainage apparatus evolved to serve as conduits for tear flow. Little is known about the ontogenesis of this system. Here, we investigated tear duct origin, developmental course, genetic and cellular determinants in mouse. We report that primordial tear duct (PTD) originates from junction epithelium of the joining maxillary and lateral nasal processes, which reshapes into future tear duct branches. We identified Prickle 1 as a hallmark for tear duct outgrowth, ablation of which stalled duct elongation. In particular, the disruption of basement membrane (BM) with cytoplasmic accumulation of laminin suggests aberrant protein trafficking. Mutant embryoid bodies (EBs) derived from iPSCs recapitulate BM phenotype of the PTD exhibiting defective visceral endoderm (VE), which normally expresses high level of Prickle 1. Furthermore, replenishing mutant VE with Prickle 1 completely rescued BM but not cell polarity. Taken together, our results reveal a distinct role of Prickle 1 in regulating polarized BM secretion and deposition in precedently uncharacterized tear drainage system and VE, which is independent of apicobasal polarity establishment.

## Introduction

A main function of tear flow is to lubricate ocular surface preventing it from drying. Disturbing tear production or flow would lead to unhealthy ocular surface including dry eye syndrome, which has a global prevalence ranging from 5%∼ 50% ^1^. In many tetrapods, orbital glands and excretory/drainage conduits coevolved to produce and passage tears over the ocular surface ^2^. The drainage conduits consisting of canaliculi (CL) and NLD (nasolacrimal duct) canals (Burling et al., 1991; Thiessen D, 1992) also have absorptive and secretory functions, possibly to provide feedback in tear production ^3-5^. Despite its importance, the drainage system remains largely unexplored.

A few comparative studies demonstrated considerable phylogenetic variations in drainage ducts ^2, 4, 6-10^. The human NLD is of ectodermal origin and starts to develop at about 5.5 weeks of gestation ^11^. Epithelial maxillary and nasal processes form a lacrimal groove, which then pinches off as a solid cord extending toward both orbital and nasal directions ^11^. The rabbit NLD shares many similar anatomical aspects with that of humans and is considered a suitable model for human NLD study ^2, 4^. However, unlike human, the rabbit NLD originates at the subcutaneous region of the lower eyelid extending unidirectionally to the naris ^7^. Similar observation was reported in a rodent animal, Mongolian gerbil ^8^. Mouse NLD is assumed to develop similarly as humans based on a scanning electron microscopy study (Tamarin and Boyde, 1977). However, such a notion faces challenges in that both the mouse and Mongolia gerbil are taxonomically of Rodentia, evolutionarily much closer to rabbits than humans. Thus, an examination of ontogeny of the mouse NLD shall resolve such contradiction.

To date, developmental and genetic determinants of the tear duct are still lacking. CL and NLD defects are seen in several syndromic diseases with known causal mutations ^12-18^. Among those, mutations in FGF signaling components (FGF10, FGFR2 and FGFR3) cause hypoplasia of NLD and lacrimal puncta often with conjunctivitis as part of Lacrimo-auriculo-dento-digital (LADD) syndrome ^18^. Consistent with these findings, mutations in p63, upstream of FGF signaling, leads to disorders overlapping with LADD, in which lacrimal outflow obstruction can occur with agenesis of NLD and CL ^19^. With congenital nasolacrimal duct obstruction (CNLDO) being present in up to 20% of newborn infants ^20, 21^, it is conceivable that many more genetic and/or genetic risk factors being involved in the tear duct obstruction and development.

The Wnt/PCP pathway plays key roles in morphogenesis of diverse tissues during development^22-24^. Particularly for tubulogenesis, tubular branching, elongation and migration require PCP-driven convergent extension (CE) and oriented cell division ^25, 26^. A set of six proteins including Prickle, Frizzled, Disheveled, Vangl, Diego, and Flamingo executes Wnt/PCP signaling to control cell polarity and oriented cell migration ^27, 28^. Mutations in *Vangle, Frizzled, Prickle* and a non-core PCP component, protocadherin *Fat*, all cause malformed renal tubules ^25, 29, 30^. Because tear duct development is a process of tubulogenesis, we hypothesized that it would require Wnt/PCP signaling, as demonstrated in other tubular organs. To test our hypothesis, we investigated the full course of tear duct ontogenesis in mouse and identified a crucial role of Prickle 1 in tear duct elongation. We further show a general role of Prickle 1 in regulation of polarized secretion and deposition of basement membrane (BM) in tear drainage system and embryoid body (EB) organoids, which is likely independent of establishment of apicobasal polarity.

## Results

### Tear duct origin and the timings of crucial events during development

To gain developmental insights into tear drainage system, we first determined when tear duct initiates by performing 3D-reconstruction of primordial tear duct (PTD) at E11 using iDISCO technology ^31^ (Fig. 1A, Supplemental movie 1). From different perspectives, tear duct was seen to initiate from the epithelial junction of fusing maxillary and nasal plate ectoderm (Fig. 1A) to form an epithelial cord. This was likely achieved partially through epithelial-mesenchymal transition (EMT), which generated a cell mass with multipolar protrusions observed on sections (Fig.1B-E, F-I). A thinning stalk connecting with maxillary/nasal surface ectoderm was detected at E12 (Fig. 1F-I). The anterior/nasal portion of the PTD had already extended a considerable distance at this age but could not be visualized together with orbital PTD on same section (see later).

**Figure 1.**
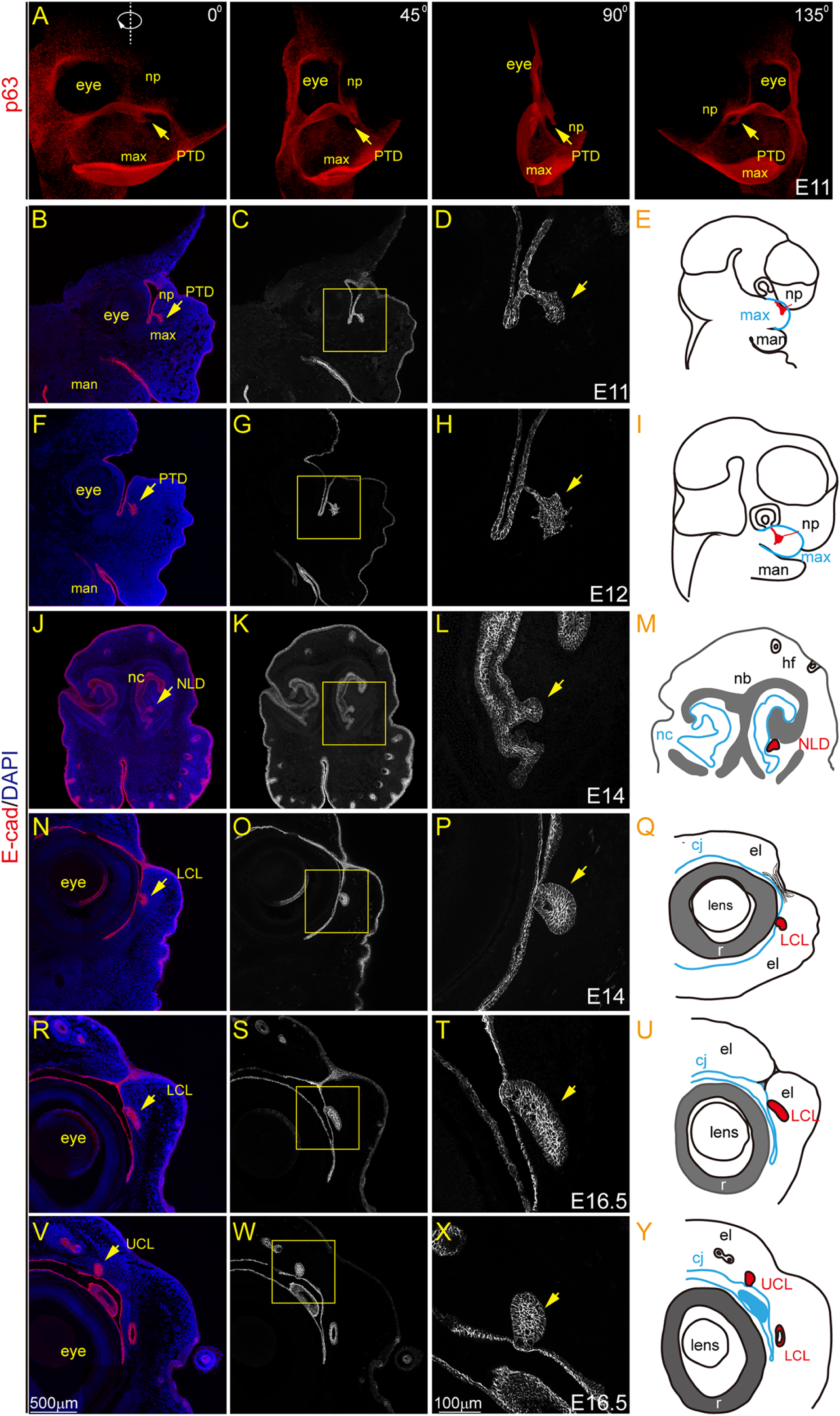
Tear duct origin and the timing of completion of the drainage system development. (A) Still images from 3D-reconstruction of p63-stained E11 head images using iDISCO. Sub panels are different view angles as indicated in up corners of the images. Arrows indicate primordial tear duct (PTD). max, maxillary, np, nasal process. (B-Y) All sections were stained with E-cadherin. Boxed areas are magnified to the right. (B-D) A parasagittal section of E11 mouse head showing initiation of primordial tear duct (PTD) from the fusing epithelium of maxillary (max) and nasal processes (np) (arrows). (E) Schematic illustration of a mouse head with the position where the PTD (red) was born. man, mandibular process. (F-I), A E12 parasagittal section showing epithelial cells transform in shapes and polarity. (J-M) A frontal section at E14 demonstrating nasolacrimal duct (NLD, arrows) fusing with nasal cavity (nc) epithelium (arrows). nb, nasal bone; hf, hair follicles (N-Q) Lower canaliculus (LCL, arrows) fusing with the lower eyelid conjunctiva (cj) at E14 on a parasagittal section. (R-U) LCL fusing with lower eyelid conjunctiva at E16.5. (V-Y) Upper canaliculus (UCL, arrows) fusing with the upper eyelid. el, eyelid.

To find the timing of PTD reaching target tissues, we first sectioned mouse head coronally to determine when NLD reaches the nasal cavity. NLD extended close to the nasal cavity but did not reach it at E13.5 (Supplemental Fig. 1A-F). At E14, NLD reached the nasal cavity and merged with nasal epithelium (Fig. 1J-M). Because the positioned mouse head for sectioning was not completely symmetrical to vertical axis, only one side of NLD was shown. The opposite side was beyond the joining point of NLD and the nasal epithelium. We next prepared parasagittal sections to define the timing of canaliculi joining the conjunctiva. The lower CL (LCL) started to join conjunctival epithelium at E14 (Fig. 1N-Q), when the upper CL (UCL) still had a distance to reach the upper eyelid (Supplemental Fig. 1G-I). By E16.5, both LCL and UCL joined the conjunctival epithelium of the lower (Fig. 1R-U) and the upper eyelid (Fig. 1V-Y), respectively. Thus, the data together demonstrated that the tear duct originated from maxillary/nasal epithelium starting at around E11 and completely reached target tissues by E16.5.

### *Prickle 1* is a hallmark for tear duct ontogenesis

We next investigated whether Wnt/PCP signaling components, which play essential roles in tubulogenesis in a variety of tissue contexts, were expressed in PTD using in situ hybridization. A number of Wnt/PCP genes were expressed in developing tear duct (not shown). Among those, *Prickle 1* is expressed exclusively in PTD but not conjunctival epithelium. Taking advantage of a knock-in eYFP reporter in *Prickle 1* heterozygous null allele^32^, we examined *Prickle 1* expression at a series of embryonic ages. *Prickle 1* was found strongly expressed in initiating/primordial tear duct at E11 on parasagittal sections (Fig. 2A-C, green). On horizontal sections, *Prickle 1* was also seen to be strongly expressed at the frontal area of maxillary process, and moderately but specifically in initiating tear duct (Fig.2D, E). Similar expression was found at E11.5 and E12 parasagittal and horizontal sections, when the tear duct continued growing out and the orbital/posterior branched out (Fig. 2F-O). Bifurcation of canaliculi (CL) and anterior NLD were clearly seen at E13 on separate parasagittal sections strongly expressing *Prickle 1* (Fig. 2P-U), which continued through until CL and NLD reaching target tissues (Supplemental Fig. 2A, frontal, B-D, parasagittal). The data thus defined *Prickle 1* as a bona fide marker for tear duct ontogenesis.

**Figure 2.**
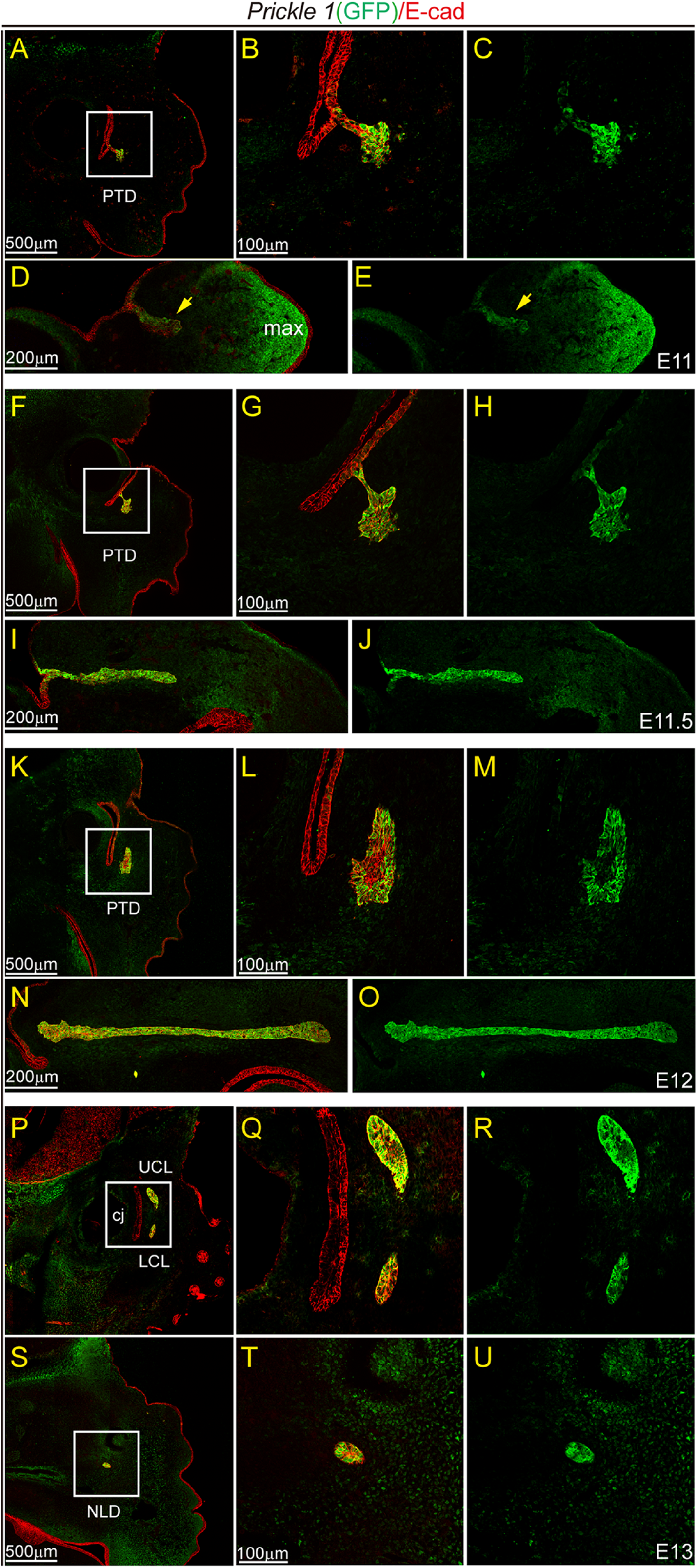
Expression of *Prickle 1* during tear duct development. In all panels, *Prickle 1* expression is indicated by a genetically knock-in *eYFP* reporter stained with anti-GFP antibody (green). E-cadherin stains general epithelium (red). Boxed areas are magnified in right panels of the same row. (A-C) A parasagittal section at E11 showing Prickle 1 expression in PTD initiating budding epithelial cells. Boxed area in (A) is magnified in (B) (merged) and (C) (separate Prickle 1/GFP channel). (D, E) Horizontal sections of the same age of PTD demonstrating extended anterior PTD highly expressing *Prickle 1*. (D), merged image. (E), separate Prickle 1/GFP channel. (F-J), Tissue sections at E11.5 with same arrangement as in (A-E). (K-O), Tissue sections at E12 with same arrangement as in (A-E) and (F-J). (P-U), Parasagittal sections at E13 demonstrating UCL and LCL branches near to the orbital conjunctiva (P-R) and advancing NLD (S-U).

### Disruption of *Prickle 1* stalled tear duct outgrowth

The unique *Prickle 1* expression pattern prompted us to query its function in tear duct development. Using *Prickle 1* null mutants that were created previously ^30^, we first performed 3D-reconstruction of serial age progression of TD from both wild type and mutant embryos from E11 to E14 at half-day intervals. This offered us an additional opportunity to have an overall look at full course of tear duct development. Consistent with the previous observations (Fig. 1 & 2), on lateral views, wild type tear duct anlage/primordial tear duct (PTD) was detected as early as embryonic day 11 (E11) (Fig. 3A). As the PTD elongated, the anterior became thinner and advanced nasally, whereas the posterior was thickened with a narrow stalk connected to the original surface ectoderm (Fig. 3B, C). By E12.5, PTD completely separated from the surface ectoderm and was transformed into three tubular branches (Fig. 3D). The long anterior branch was the future nasolacrimal duct (NLD), and the shorter posterior branches were prospective upper and lower canaliculus (UCL and LCL), respectively. The three tubules continued growing toward their prospective targets—the conjunctival and nasal epithelia (Fig. 3E-G). The whole tear drainage system -- the canaliculi and NLD, and its positional relationship to conjunctiva and nasal cavity was clearly recognized after E12.5 (Fig. 3E-G).

**Figure 3.**
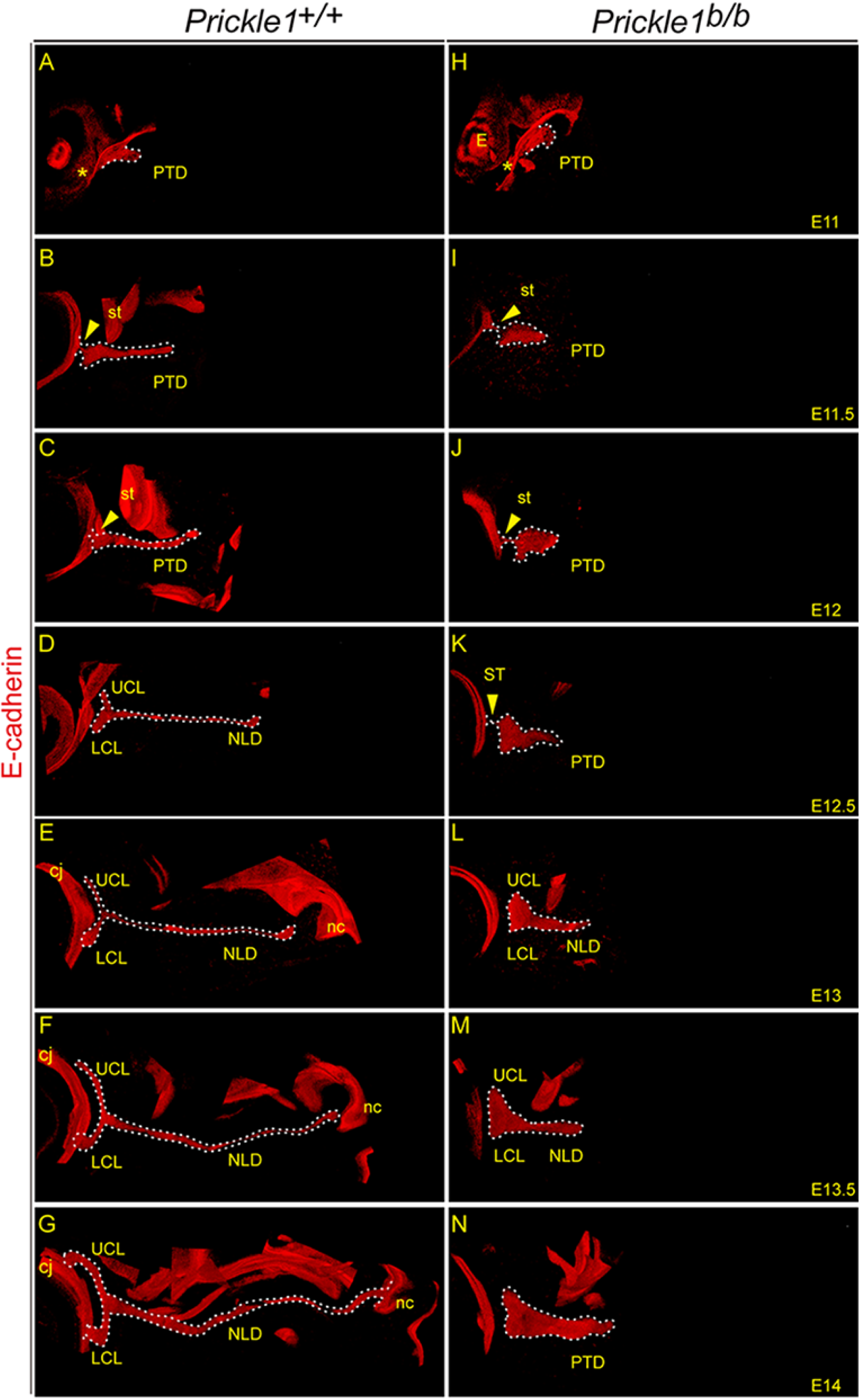
Disruption of *Prickle 1* stalled tear duct outgrowth. E-cadherin staining was used to identify developmental tear duct in all panels. Vibratome sections at 100 μm were used for collecting stacks of images for 3D-reconstruction. Only side view was shown in the figure. (A-G), Developing wild type tear duct system at serial embryonic ages. Note the connecting stalk (arrows) at E11.5 (B) and E12 (C), and the branching for prospective canaliculi at E12.5 (D). (H-N), Mutant tear duct at comparable ages of the wild type shows shortened and broadened features. Arrows point to the connecting stalks before E13.

The mutant PTD was initiated at about the same time as the wild type (Fig. 3H), however, it was shorter and wider at all ages examined (Fig. 3I-N). Separation of the epithelial stalk from the presumptive conjunctiva was accomplished at E13 (Fig. 3L), half a day later than that of the wild type, which was complete at E12.5 (Fig. 3D). Collective movement/migration of the outgrowing epithelial cells was hampered (Fig. 3I-N), and characteristics of canaliculi and NLD branches were barely identifiable in the mutants (Fig. 3K-N). Thus, ablation of *Prickle 1* stalled tear duct outgrowth, which is consistent with its expression in this organ.

### Disruption of *Prickle 1* impaired cell-cell adhesion and polarized BM secretion

To examine cellular changes in the mutant PTD, we first examined E-cadherin/catenin junction complexes on horizontal sections of E11.5 embryos. β-catenin and E-cadherin were normally colocalized at the cell junction throughout PTD (Supplemental Fig. 3, C, D, G, H, Fig. 4C-D). In the mutants, however, β-catenin mostly translocated to the cytoplasm, and E-cadherin staining was patchy rather than continuous (Fig. 4E-G, Supplemental Fig. 3E, F, I, J). Of note, proximal mutant PTD often had relatively normal junctional localization in comparison to other areas (Fig. 4 E-G). Further examination of α- and p120 catenins revealed a regional difference for their localization of in that the advancing tip in wild type mice exhibited more cytoplasmic staining than rest of the tube (Supplemental Fig. 3K, L, S, T). This regional difference also existed in basement membrane components (see later). Regardless, like β-catenin, both α- and p120 catenins were ectopically located in cytoplasm throughout the mutant duct along with patchy E-cadherin in majority of areas (Supplemental Fig. 3M, N, Q, R, U, V, Y, Z). Additionally, the broadened and shortened mutant PTD appeared to have massive loosely connected cells in the presumptive lumen (Fig. 4E-G, Supplemental Fig. 3, and later).

**Figure 4.**
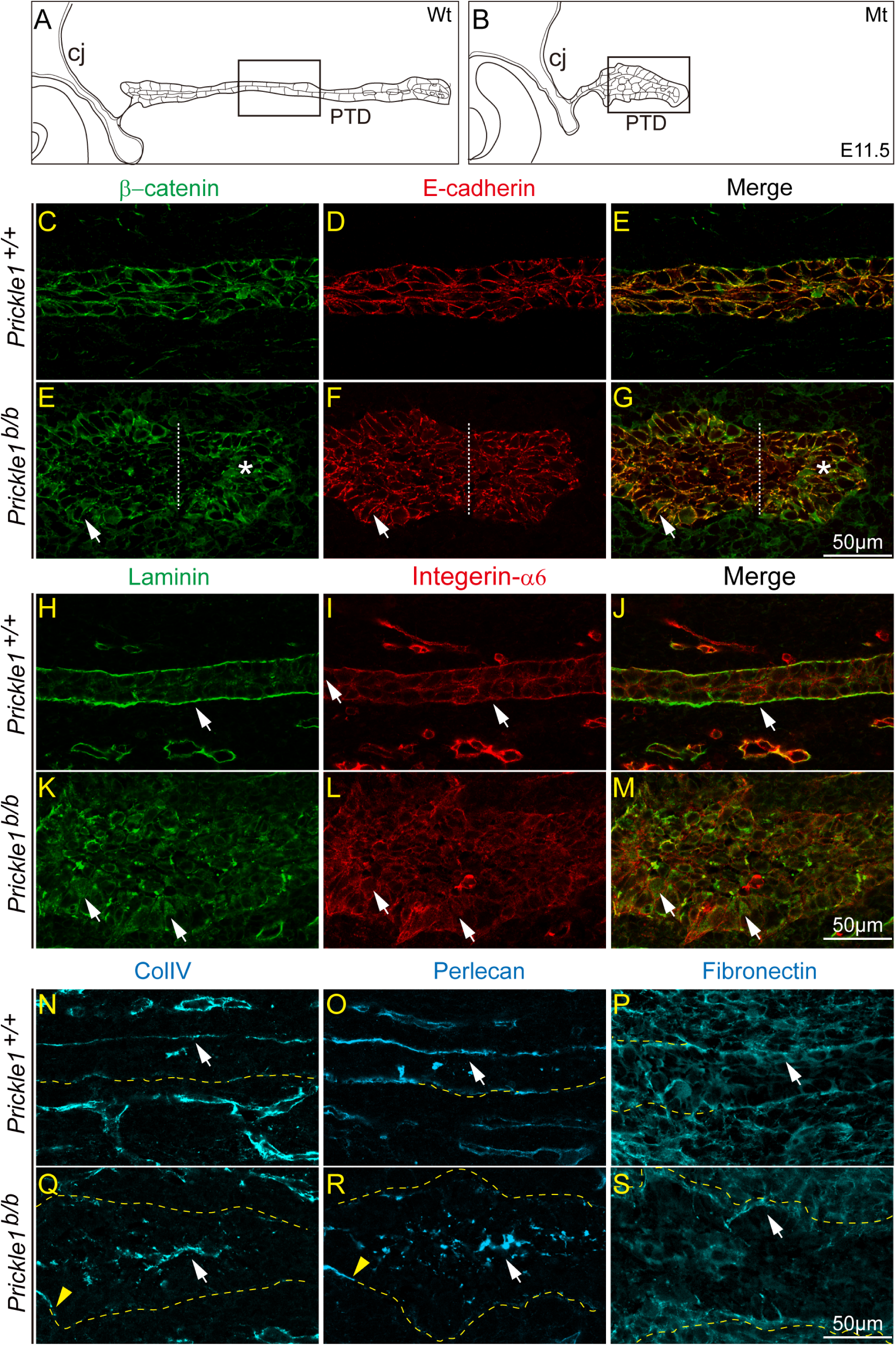
Disruption of *Prickle 1* impaired cell-cell adhesion and cell-matrix adhesion. (A-B), Schematic illustration of developing tear duct at E11.5. (A), Wild type tear duct (Wt). (B), Mutant tear duct (Mt). Boxed areas are roughly corresponding to the regions demonstrated in below panels respective for wild type and the mutant duct. (C-G), Tissue sections co-labeled with β-catenin (green) and E-cadherin (red). Dashed lines roughly divide anterior/orbital and posterior/nasal halves of PTD in the mutants. Arrows within posterior halves of PTD in (E-G) indicate relative normal localization of β-catenin and E-cadherin in the mutants, whereas arrows within anterior halves indicate cytoplasmic β-catenin localization. (H-M), Tissue sections co-labeled with Laminin (green) and Integrin-α6 (red). Arrows in wild type tube (H-J) point to basal deposition of Laminin and Integrin-α6 receptor, while in the mutants indicate cytoplasmic localization of Laminin and Integrin-α6 (K-M). (N, Q), Col. lV staining. Col. lV basal deposition in wild type tube (N, white arrow) becomes mostly luminal in the mutants (Q, white arrow). Yellow arrowhead point to the initial part of the mutant PTD having normally deposited BM. (O, R), Perlecan staining. Same deposition pattern is seen as Col. lV in both wild type and the mutants. (P, S). Relatively normal basal deposition of Fibronectin in both wild type (P) and the mutants (S). Yellow dashed lines drawn to assist to visualize PTD shape.

The accumulation of luminal cells in the mutant PTD suggested defective cell-matrix adhesion, which could be simply due to comprised BM. An examination of extracellular matrix (ECM) proteins of the BM confirmed this speculation. In wild type, BM components were deposited to basal lamina (Fig. 4 H, J, N, O) and displayed a notable loss in the advancing tip area of the PTD (Supplemental Fig. 4D, J, P). In the mutant, laminin, Col.IV, and Perlecan were all mislocalized and BM was disrupted in all areas but the initial stalk (Fig. 4K, M, Q, R). Considerable amount of laminin was trapped in the mutant cytoplasm along with Integrin α-6 receptor (Fig. 4K-M). Col.IV and Perlecan were instead secreted ectopically in the presumptive lumen/apical side (Fig. 4 Q, R). Interestingly, basal localization of fibronectin was largely preserved throughout the mutant PTD (Fig. 4P, S). Taken together, both cell-cell and cell-matrix adhesions were disrupted in the mutant PTD.

### Disruption of *Prickle 1* altered cytoskeleton and vesicle compartments

The endoplasm-trapped and ectopically secreted BM components in the mutant PTD cells led us to further examine cytoskeletal organization, which is directly involved in secretory pathways. The massively disorganized mutant luminal cells prevented us from gathering comparable information with that of wild type, thus we only examined the duct wall cells. Heavily stained actin bundles were observed in both wild type and mutant PTD at E11.5 (Fig. 5A-D). However, actin filaments in the mutants appeared less packed and often formed spikes crossing tissue boundary (Fig. 5B, D). Apical actin fibers overlying horizontally on the free lumen surface was often found in wild type PTD (Fig. 5A, C), but rarely distinguishable in the mutant (Fig. 5B). Similarly, the horizontally run stable microtubule tracks in wild type labeled with acetylated alpha tubulin were not detected in the mutants (Fig. 5E-H).

**Figure 5.**
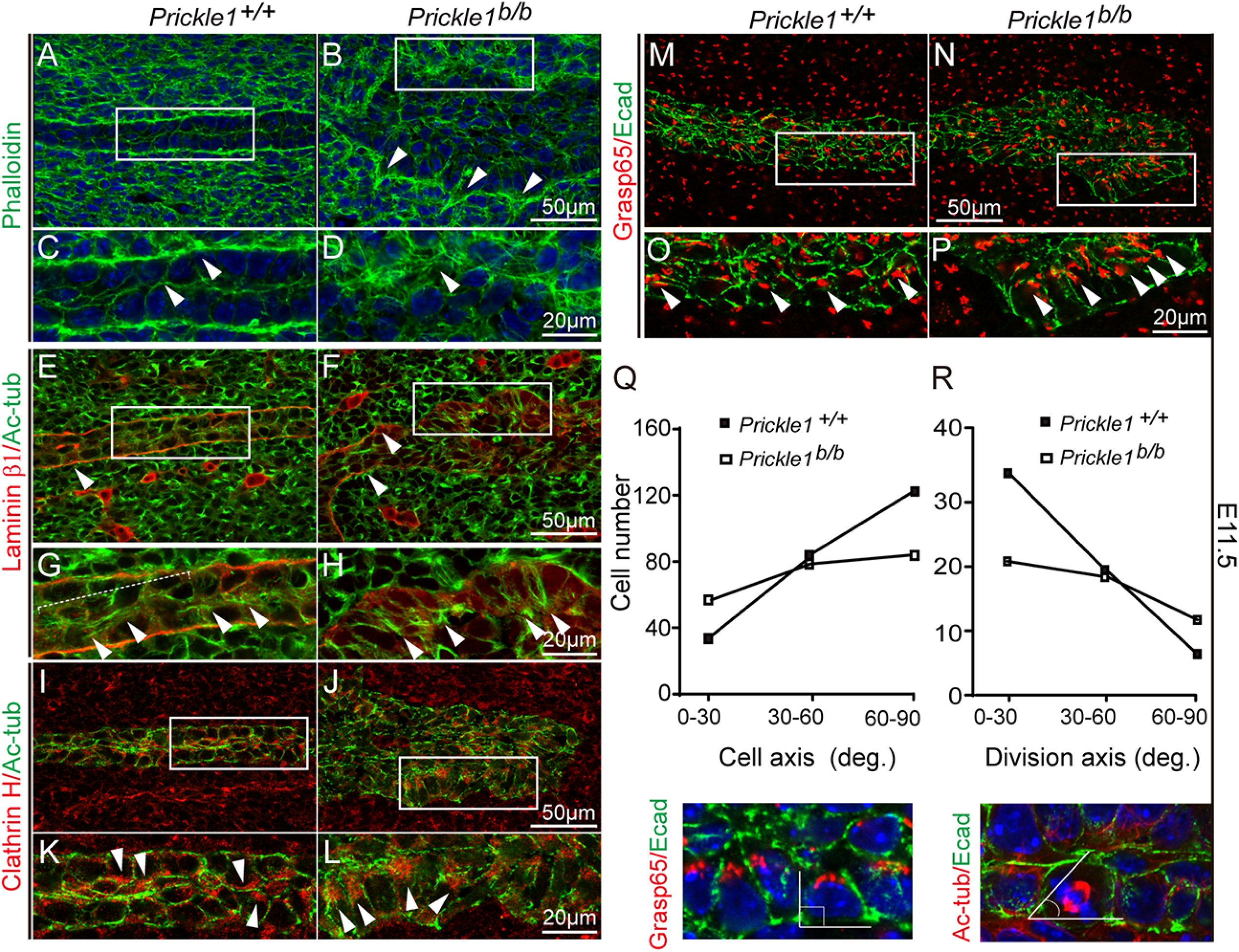
Disruption of *Prickle 1* altered cytoskeleton and vesicle compartments. Each boxed area is magnified in the corresponding below panel. (A-D), Phalloidin staining of actin filaments. Arrowheads point to basal and/or apical actin bundles. (E-H) Acetylated α-tubulin (Ac-tub, green) and Laminin β1 (red) staining. Arrowheads in (E) and (F) point to basement membrane labeled by Laminin β1 (red). Arrowheads in (G) and (H) point to orientated microtubule tracks. (I-L), Clathrin and E-cadherin staining. Arrowheads point to vesicle distributed locations. (M-N) Grasp 65 staining. Arrowheads point to apically stained cis Golgi cisternae. (Q), Quantification of cell axis orientation using Grasp 65 and E-cadherin to define apicobasal axis. (R), Quantification of cell division orientation using acetylated α-tubulin and E-cadherin to define division angle.

Because vesicle trafficking on microtubules and actin fibers directs proteins to different pathways, we examined whether endoplasmic compartments in the mutant PTD were also affected. By examining endocytic vesicles, the majority of clathrin-coated endosomes were localized along the apical domain of the wild type PTD (Fig. 5I, K), but were considerably expanded along the apicobasal axis in the mutant (Fig. 5J, L). In contrast, Golgi complex appeared grossly normal in the mutants in terms of localization (Fig. 5N, P compared to M, O). We further measured the cell axis and division orientations, which are key features of PCP, referring to the direction of PTD extension. Axis and division orientations indicated by Golgi complex and acetylated tubulin, respectively, were both randomized in the mutants (Fig. 5Q, R). These data were consistent with the observed protein trafficking defect of BM components in the *Prickle 1* mutant.

### iPSC-derived EBs recapitulated BM phenotypes of the mutant PTD

To obtain further molecular insights into the BM and cellular changes of the *Prickle 1* mutant PTD, we generated iPS cells by the “4-factor” reprogramming of mouse embryonic fibroblasts ^33^ (Supplemental Fig. 5A). iPSCs were first verified molecularly (Supplemental Fig. 5A-K), then functionally (Supplemental Fig. 5, L-N), and further differentiated into EBs ^34^ (Fig. 6A). Under bright field microscopy, the majority of EBs (Fig. 6B, 85% for control and 67% for mutant) displayed visceral endodermal (VE) cells with high *Prickle 1* expression (Fig. 6C, E, F, G, I, J). However, the mutant VE were flattened compared with that of the control (Supplemental Fig. 6A-F). The results suggested that sorting of VE layer was successful, but cell polarity may have changed.

**Figure 6.**
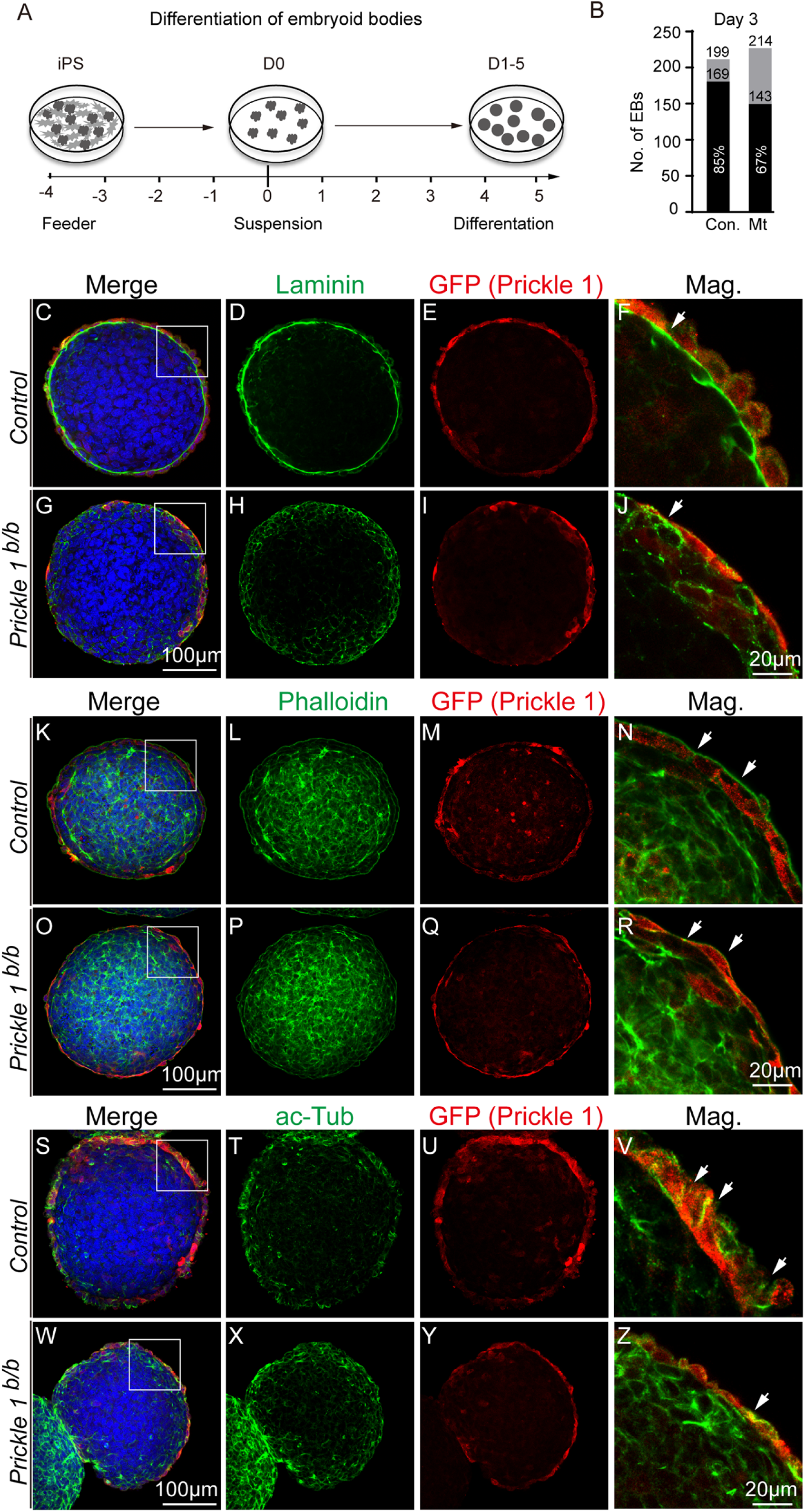
iPSC-derived EBs recapitulated BM phenotypes of the mutant PTD. (A), Schematic illustration of differentiation of EBs from iPS cells. (B), EBs differentiation efficiency. Successful EB differentiation was defined by existence of visceral endodermal (VE) cells, which express *Prickle 1*. (C-Z), Boxed areas were magnified on right most column. Red channel for all panels is staining with antibody against GFP, which reports endogenous *Prickle 1* expression. DAPI (blue) stains nucleus. (C-J), Laminin (green) staining BM, arrows point to Laminin localization. (K-R), Phalloidin staining (green), arrows point to cortical actin. (S-Z), Acetylated α-tubulin staining (green), arrows point to microtubule tracks.

We next examined the BM deposition and cytoskeleton changes in differentiated EBs. Control EBs possessed uniform BM underlying the columnar epithelial sheet as indicated by laminin staining (Fig. 6C, D, F). However, the mutant EBs failed to form basement membrane (Fig. 6G, H, J). The loss of BM was further confirmed by mislocalization of Col. IV and Perlecan (Supplemental Fig. 7). Phalloidin-stained actin filaments clearly outlined apical surface of the control endodermal cells (Fig. 6K, L, N). In the mutant, actin bundles were still present but thinner, without obvious asymmetrical distribution (Fig. 6O, P, R). Microtubule tracks labeled by acetylated tubulin were roughly parallel to apicobasal axis in control VE cells (Fig. 6S, T, V), but poorly organized in the mutants (Fig. 6W, X, Z). We further tested, at extreme conditions, whether disruption of actin filaments and microtubule tracks would alter BM and Laminin secretion. Normal EBs treated with Cytochalasin D displayed cytoplasmic retention of Laminin until 10 hr post treatment, with substantially retained BM (Supplemental Fig. 8A-H, Q). In contrast, disruption of microtubule networks with Nocodazle treatment acted much faster on breaking BM than disruption of actin filaments, typically within a 2 hr window of treatment (Supplemental Fig. 8I-P, R). While these results can’t be interpreted simply due to effects of cytoskeleton on many cellular activities, they generally agreed with the abnormal BM secretion and the altered actin and microtubule network in the mutant EBs and PTD.

### Rescue of BM phenotype but not AB polarity of the mutant EBs by expressing *Prickle 1*

The severe BM and cell polarity phenotypes in both mutant PTD and EBs implied a conserved role of Prickle 1 in these molecular events. We next sought to rescue BM and/or cell polarity by adding back *Prickle 1* to mutant EBs. Using a lentiviral system with built-in tetracycline inducible components, we infected mutant iPS cells to control *Prickle 1* expression by addition of doxycycline and harvested EBs at different time points (Fig. 7A-C). The mutant EBs showed rescue of BM starting from differentiation D2 and continuing to increase by D5 (Fig. 7C). Nearly 60% (Fig. 7C) of mutant EBs were rescued with integral BM upon *Prickle 1* induction (Fig. 7F, G, H), in contrast to less than 10% EBs having BM in control groups with either induced *Cherry* or uninduced *Prickle 1* (Fig. 7D, E). Surprisingly, although the mutant outer layer endodermal cells expressing *Prickle 1* were generally sparse (Fig. 7F, G, H), apparently not forming lateral junctions and apicobasal polarity, yet they demonstrated a robust rescue of BM. To further confirm this observation, we immunostained EBs with E-cadherin and Rab9, a GTPase involved in processes of vescicle trafficking. As expected, no lateral junction was observed for Prickle 1 expressing cells (Fig. 8A-A”, B-B”), and normally apically localized of Rab9 was randomized (Fig. 8C,C”, D-D”). These results suggested that formation of BM and cell polarity controlled by Prickle 1 might be independent or of sequential processes.

**Figure 7.**
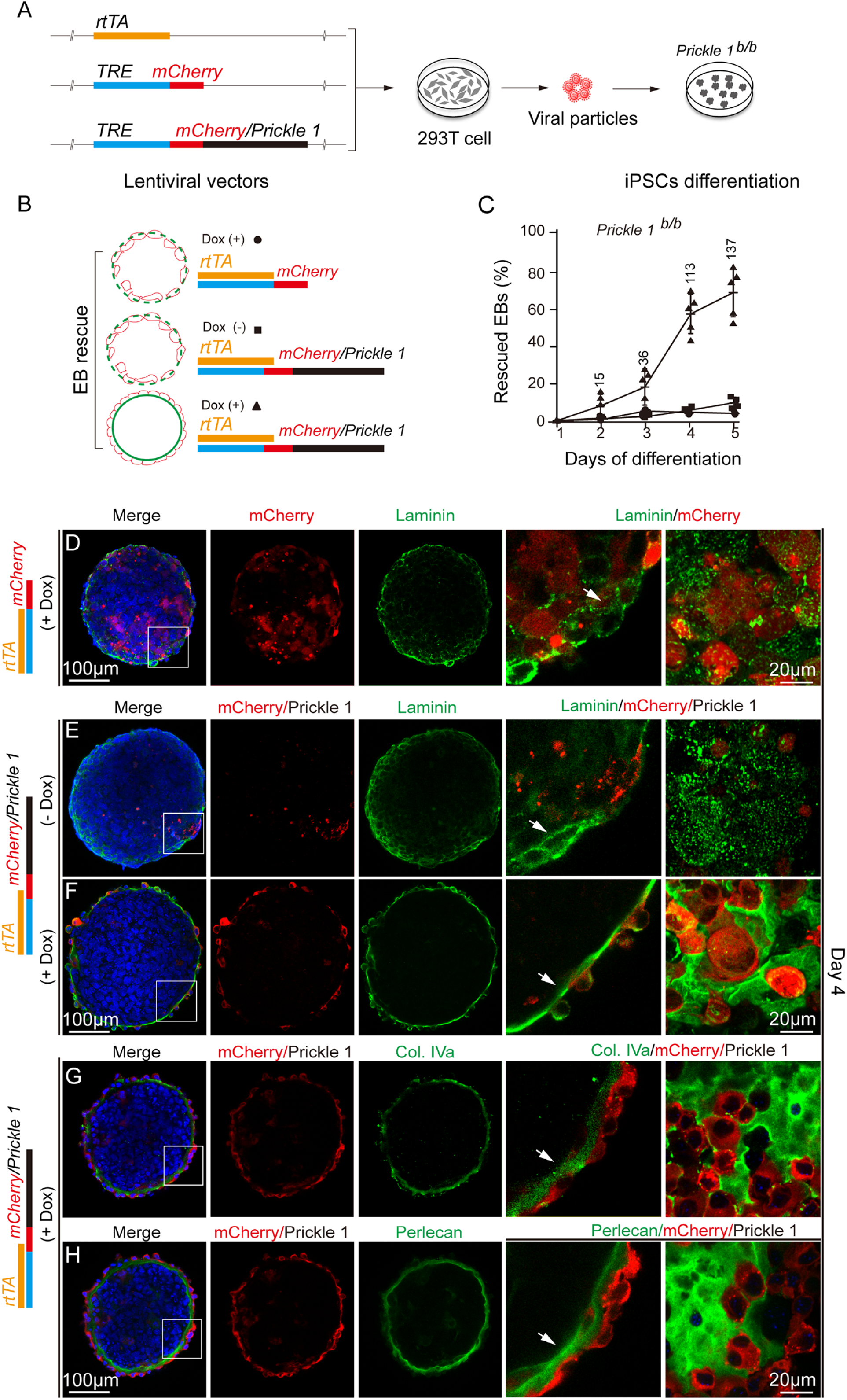
Rescue of the mutant BM phenotype in EBs by expressing *Prickle 1*. (A), Lentiviral tet-inducible vectors and virus production in HEK 293 cells. Lentiviral vectors of *rtTA, Cherry* and *Cherry/Prickle 1*-fusion genes. *Cherry* and *Cherry/Prickle 1* are under control of tet-responsive element. Vectors were transfected into HEK 293 cells together with helper viral genes for packaging. (B). Schematic illustration of BM rescue experiment for *Prickle 1* mutant EBs. *Prickle 1* mutant EBs with induced *Cherry* (Dox +) or uninduced (Dox -) *Cherry/Prickle 1*-fusion genes serve as controls for induced *Cherry/Prickle 1* expressing EBs. (C), BM rescue rate under controls and *Prickle 1* expressing conditions in mutant EBs. (D-H), Boxed areas in the first column of panels were magnified in the column next to the last. The last column of images is superficial view of the immuostained EBs. Each row of panels is from a same EB expressing *Cherry* (D) or *Cherry/Prickle 1*-fusion gene (E-H) and stained with an indicated BM marker. Arrows point to BM locations. (D), Induction of *Cherry* expression does not rescue mutant BM. (E), Without Dox, *Cherry/Prickle 1* is minimally expressed (red) without BM rescue. (F), With Dox (Dox +), sparse Prickle 1-positive VE cells lining the surface completely rescue BM deposition of Laminin. (G, H). With Dox, BM deposition of Col. IV and Perlecan were rescued.

**Figure 8.**
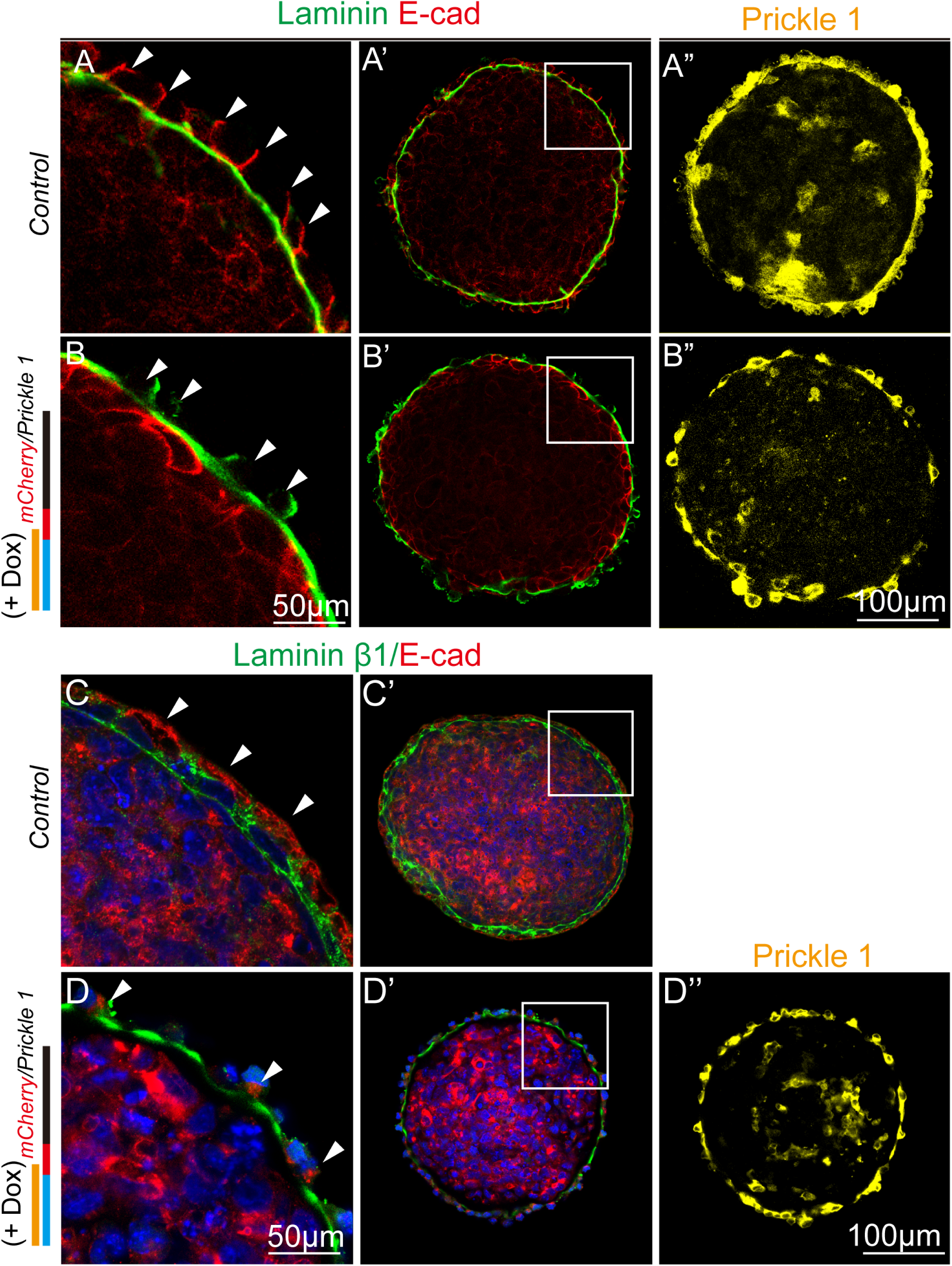
Rescue of BM in the mutant EBs is independent of AB polarity. (A, A’ A’’), A control EB labeled with E-cad (red), laminin (green) and GFP (yellow, anti-EGFP staining). (A), Magnified boxed area in (A’). Arrows point to lateral junctions. (A’) A full view of stained EB. (A’’) Prickle 1 expression revealed by eYFP reporter (anti-EGFP). (B, B’, B”), A rescued mutant EB by replenishing Prickle 1 expression. Same arrangement of panels as in (A, A’ A’’). (B) and (B’) are labeled with E-cad (red) and laminin (green). (B”), mCherry fluorescence tag of Prickle 1. Arrows point to VE cell bodies. (C, C’), A control EB labeled with Rab9 (red) and laminin β1 (green). (C), Arrows point to Rab9 staining from boxed area in (C’). (C’), A full view of stained EB. Note that Prickle 1 expression is not shown due to conflict usage of antibody species. (D, D’), A rescued EB with same arrangment of panels and staining as in (C, C’). (D”), mCherry fluorescence tag of Prickle 1.

## Discussion

Scant attention has been paid to the tear drainage system despite its importance. A few studies primarily focused on comparative anatomy between species ^2, 4, 7, 11^. The current study details ontogenesis of the mouse tear drainage system and outlines several crucial events - tear duct origin, elongation, and reaching final destinations. The study further offers genetic and molecular insights into tear duct tubulogenesis, highlighting polarized BM secretion controlled by Prickle 1 as a crucial event for duct elongation.

The formation of mouse tear duct is distinct from all other studied animals based on the current study. Origin wise, the mouse tear duct is more like that of humans, both of which initiate from joining maxillary and nasal process ^11^. However, initiation of the mouse PTD is continuous with orbital conjunctival epithelium rather than the nasolacrimal groove in humans^11^. Mouse tear duct appears to share a common origin with conjunctiva, as both are from joining surface ectoderm of maxillary and nasal plates. Fusion of the two is a zipping process from frontal/nasal to orbital/rare, leaving an orbital epithelial notch for PTD outgrowth. As the PTD separates from the conjunctival epithelium, the fusion is accomplished. The CL branches later reconnect to conjunctiva to serve as drainage conduits. Although mouse and Mongolia gerbil are both taxonomically Rodentia, origin of tear duct in the gerbil is more similar to the one of the rabbit, which is from the subcutaneous region of the orbital lower eyelid ^7, 8^. The differences of tear duct genesis in different species likely reflect craniofacial features adapted for survival needs.

The expression pattern of Prickle 1 in tear duct is unique. Unlike many other genes we identified, most of which are expressed in both conjunctiva and PTD (not included), *Prickle 1* is found in budding tear duct excluding conjunctival epithelium. This pattern emphasizes a specific need for *Prickle 1* during tear duct development. From initiation to reaching final destinations, the ductal cells maintain epithelial fate (as indicated by E-cadherin expression), but undergo dynamic depolarization, repolarization and intercalation. As a PCP signaling component, *Prickle 1* may serve most of these needs since disruption of *Prickle 1* altered cell polarity, intercalation and division orientation. It would be intriguing to know how spatial and temporal expression of *Prickle 1* is determined in tear duct initiation. As a general principle in developmental biology, positional inductive cues (likely from the fusion of maxillary and nasal processes in this case) is probably involved in *Prickle 1* activation, which needs to be further investigated.

Striking cellular changes in *Prickle 1* mutant PTD are disrupted BM deposition and cell-cell adhesion. Basal deposition of BM does not always coincide with adherens junction formation as observed from both wild type and the mutant PTD. The data thus suggests that regulation of Prickle 1 function in BM might be independent of apical cell junctions. How then is Prickle 1 involved in BM deposition and cell adhesion?

The cytoplasmic accumulation of laminin and ectopic deposition of other BM components indicate a misrouted secretory pathway(s). However, the impaired trafficking does not appear generalized to all proteins because basal deposition of fibronectin is largely retained in the mutant wall cells, and even adherens junctions are not uniformly disrupted. Additionally, coordinated polarization of Golgi complex and Clathrin-coated vesicles is largely unaffected. Together with altered mutant actin filaments and microtubule tracks, the data suggest that trafficking of a select set of proteins (including BM) are affected by loss of *Prickle 1*. Of note, a recent study in Drosophila reported that *Prickle* also modulates microtubule polarity impacting axonal vesicle transport ^35^. Whether vesicle trafficking in epithelial cells and neurons is similarly regulated by Prickle 1 remains of interest to be unveiled.

The crucial role of Prickle 1 in BM deposition and polarity in two tissue types--PTD and EB, hints at a converging role of Prickle 1 in BM formation. If so, restoring *Prickle 1* expression alone would rescue both BM and polarity in these tissues. Yet, adding back Prickle 1 to the mutant EBs only rescued BM, with sparse VE cells without basolateral domain or apicobasal polarity. While it is possible that the restored amount of Prickle 1 might be inadequate for polarity rescue, the result does imply that Prickle 1-regulated BM either precedes or is largely independent of general polarity machineries. Another finding from the EB study is that the visceral endoderm layer (VE) highly expressing *Prickle 1* is crucial for formation of BM between VE and inner cell mass. These findings contrast with the previous suggestion that ECM deposits according to AB polarity of the epiblast ^36^. Thus, the VE polarity, migration and epiblast polarity might be coupled through BM, emphasizing an non-tissue-autonomous role of Prickle 1 in epithelial polarity setup.

PCP cross-talking with ECM has been demonstrated in axis elongation of multiple organisms including ascidians ^37^, fly ^38^ fish and frog ^39, 40^. Tubulogenesis driven by PCP has been well studied ^25, 26^, though not from the perspective of ECM contribution. The current study, for the first time, demonstrates a crucial role of PCP-controlled ECM secretion in tubulogenesis. Notably, the Drosophila egg chamber employs an unusual form of planar polarity with aligned basal actin bundles and BM providing “molecular corset” to direct chamber elongation ^38^. The similar process may also be utilized for PTD elongation to restrict expansive forces and promote lengthening of the duct.

In summary, our study provides novel knowledge of ontogeny of tear drainage system, which is implicated in a wide range of ocular disorders. The ontogenesis of tear duct requires Prickle 1-regulated polarized basement membrane secretion and deposition. This process is likely through a role of Prickle 1 in intracellular trafficking and secretion, which is independent of general polarity machinery. Disruption of the intrinsic polarized BM secretion will be ultimately transformed into an amplified extrinsic impact on AB or tissue polarity, which may also directly involve Prickle 1.

## Acknowledgements

We thank Dr. Tiansen Li from the National Eye Institute for providing valuable suggestions in preparation of the manuscript. The authors thank Tiansen Li, Rong Ju for critical reading the manuscript and helpful comments. We thank lab members Shujuan Xu, Shanzhen Peng, and Xinyu Gu for technical supports to the work.

## Materials and methods

### Mice and genotyping

Animal husbandry and experimentation were conducted in strict adherence to the Standards in Animal Research: Reporting of In Vivo Experiments (ARRIVE) guidelines, with approval from Animal Care and Use Committee (ACUC), Zhongshan Ophthalmic Center, Sun Yat-sen University. Mouse strains were of mixed genetic backgrounds of C57BL/6 and Sv129. *Prickle 1* gene-trap mutant strain was created as described previously ^30, 32^. Straight null allele (*Prickle 1*^*b/+*^) was created by excision of Cre recombinase driven by Sox2 promoter (Sox2-Cre)^30^. Mouse genotyping was conducted as described previously ^30, 32^. A knock-in *eYFP* reporter under the control of endogenous *Prickle 1* promoter was used to monitor *Prickle 1* expression.

### Generation of iPSCs from MEFs

Isolation, culturing, and maintenance of mouse embryonic fibroblasts (MEFs) followed standard protocols ^41^. Briefly, both *Prickle 1* wild type, heterozygous and mutant embryos were collected at post-coitus day E13.5. After removal of head and viscera, the remaining trunk was subjected to trypsin digestion and cultured in DMEM medium.

For iPS cells (iPSCs) induction, MEFs were reprogrammed into iPSCs by lentiviral particles expressing “4 factors” (Takahashi and Yamanaka, 2006) under the control of a tet-inducible expression system. Briefly, TetO-FUW-OSKM and FUW-M2rtTA (Addgene) lentiviral vectors were separately transfected into HEK 293 cells together with pMD2.G, pMDLg/pRRE and pRSV-Rev helper virus vectors (Addgene). Culture medium was collected and ultracentrifuged to concentrate viral particles. Cultured MEFs were then infected with the packaged OSKM and rtTA viruses separately, with doxycycline (SD8430, Solarbio, China) induction. iPSCs can be observed 5 days after infection. All plasmid information can be found at: https://www.addgene.org/20321/ (for TetO-FUW-OSKM), https://www.addgene.org/20342/ (for FUW-M2rtTA), https://www.addgene.org/12259/ (for pMD2.G), https://www.addgene.org/12251/ (for pMDLg/pRRE), and https://www.addgene.org/12253/(for pRSV-Rev)

The derived-iPSCs were first characterized by alkaline phosphatase staining according to the manufacturer’s instructions (MA0197, Meilunbio, China), then subjected to immunohistochemistry for examination of a set of stem cell markers including Sox2, Nanog and SSEA1 (Supplemental Fig 6). Teratoma formation was used to functionally evaluate pluripotency of iPSCs by intraperitoneal implantation of 1×10^6^ cells into immunodeficent mice ^42^. Mice were sacrificed at 2.5 months after transplantation followed by H&E staining to identify tissue types (Supplemental Fig 6).

### EB differentiation and rescue of the mutant EBs

The derived iPSCs were differentiated into embryoid bodies (EBs) according to the protocol described previously ^34^. Because heterozygous EBs carry eYFP reporter from *Prickle 1* endogenous locus facilitating visualization of the visceral endodermal (VE) layer, and were phenotypically normal as the wild type EBs, they were used as controls.

For the rescue experiment, mCherry-tagged Prickle 1 (mCherry/Prickle 1) and mCherry cDNAs were separately cloned into a lentiviral vector by replacing OSKM cassette of the TetO-FUW-OSKM vector. Virus packaging was performed by transfection of each construct into HEK 293 together with helper virus vectors as described previously. Mutant EB infection with Prickle 1 and mCherry viruses was conducted at differentiation Day 0 with addition of doxycycline. Successfully differentiated EBs can be identified with integral outer layer of visceral endoderm having *Prickle 1* expression (indicated by a knock-in EYFP reporter).

### Tissues, histology, immunohistochemistry and antibodies

Mice of postnatal day 1 were sacrificed by decapitation. Dissected mouse heads were directly embedded in OCT (SAKURA, Cat. 4583, USA), quickly frozen in liquid nitrogen, and stored at −80 °C freezer until use.

For collection of timed embryos, appearance of vaginal plug was designated as embryonic day 0.5 (E0.5). Embryos of anticipated ages were dissected out of the uterus and fixed in 4% paraformaldehyde for 24h at 4°C. Samples were washed 3 times in PBS (Phosphate buffered saline), put through a series of sucrose (10%, 20%, 30%), then embedded in OCT and stored at −80 °C until use.

For immunostaining, tissue sections were cut at 15 μm (Leica Cryostat 1900), blocked with 10% donkey serum with 0.1% Triton X-100 in PBS (PBST) for 30 min at RT, then incubated with primary antibodies at 4°C overnight. After washing with PBST, sections were incubated with fluorescent dye-conjugated second antibodies for 1 hr at room temperature, washed 3 times in PBST and mounted with Fluoromount-G (Southern Biotech, Birmingham, AL, USA).

For vibratome sections, embryos were fixed with 4% PFA, washed with PBS, then embedded in 4% agarose and sectioned with Leica VT 1000S at 100 μm thickness. For immunostaining, sections were incubated with primary antibodies for 24 hr in a 48-well culture plate with shaking. After 3 washes with PBST at 30 min each, sections were incubated with secondary antibodies overnight. Samples were washed 3 times with PBST (30 min each) and mounted on slides using Fluoromount-G.

For differentiated EBs, immunohistochemistry was performed following the same protocol used for vibratome sections.

For iDISCO ^31^ preparation, mouse tear duct at E11 was labeled with p63 antibody. Briefly, embryos were dehydrated through a series of methanol (30%, 50%, 70%, 90%, 100%) after fixation, bleached with H_2_O_2_/DMSO/methanol(1:1:4), rehydrate through methanol/PBS (80%, 60%, 40%, 20%, PBS), and treated with 2% Triton X-100. Embryos were further permeabilized following Renier et al ^31^, blocked with 6% donkey serum and incubated with primary antibody at 37°C for at least 1 day. Embryos were washed in the same incubation buffer for a day (4-5 times of changing solution), then incubated with secondary antibody and washed the same way as done for the primary.

Immunostained embryos were cleared through methanol series, then switched to Dichloromethane (DCM, 270997, Sigma), then to DiBenzyl Ether (DBE, 108014, Sigma). The embryos were ready for imaging after clearance.

Antibodies used in this study are: Anti-E-cad (ab11512, Abcam), Anti-N-cad (QF215275, Life Technologies), anti-p63 (ab124762, Abcam), anti-GFP (TP401, Torrey pines biolabs), and anti-GFP (ab6673, Abcam), anti-Laminin (L9393, Sigma), anti-Laminin β1 (ab44941, Abcam), anti-Collagen IV (ab19808, Abcam), anti-Perlecan (MA1-06821, Invitrogen), anti-Ac-Tubulin (T6793, Sigma), anti-Clatherin-H (ab172958, Abcam), anti-GRASP65 (ab30315, Abcam), 488- Phalloidin (A12379, Thermo), 568- Phalloidin (A12380, Thermo), anti-Nanog (ab80892, Abcam), Anti-Sox2 (sc365964, Santa Cruz), Anti-SSEA1(ab16285, Abcam), Anti-integrin α6 (ab105669, Abcam), Anti-integrin β4 (ab25254, Abcam), Anti-fibronectin (ab23750, Abcam), β-catenin (610153, BD), α-catenin (13-9700, ZYMED), P120-catenin (66208, Proteintech).

### Imaging and 3D-reconstruction

For P1 tear duct reconstruction, fresh frozen mouse heads were cut coronally at 30 μm, stained with anti-p63 antibody to identify tear duct on each section. For embryos from E11-14, sections were cut parasagittally at 100 μm, stained with E-cadherin antibody to identify developing tear duct.

Fluorescence microscopy images were obtained on a Zeiss confocal microscope (Zeiss LSM880, Zeiss, Oberkochen, Germany) and Imager Z2 equipped with ApoTome (Zeiss, Oberkochen, Germany). Images taken from microscopes were aligned manually by Photoshop according to anatomical features of each section. Tear duct structures were traced on each consecutive section and imported to NIH ImageJ software for 3D-processing (3D viewer plugins) with correct setting of image-depth scale, pixel depth and adjusted image coordinates.

### Drug treatment of EBs

EBs at differentiation day 2 were treated with Cytochalasin D (final concentration 10μM, PHZ1063, Thermo) and Nocodazle (final concentration 0.5μg/ml, M1404-2MG, Sigma). Vehicle (DMSO) only was used in control groups. EBs were harvested at 2 hr and 10 hr post treatment, washed with PBS for several times and fixed with PFA for further processing.

### Quantification and statistics

Cell axis orientation was quantified by angle between apicobasal axis (defined by Grasp65 staining, Fig. 6) and tubule axis indicated by E-cadherin staining. Similarly, cell division orientation was defined by angle between mitotic spindle axis indicated by acetylated α-tubulin staining and tubule axis. A total of 236 wild type and 215 mutant cells from 4 animals, and 61 wild type and 53 mutant cells from 8 animals were quantified for cell axis and cell division orientation, respectively.

